# Autonomous behaviour and the limits of human volition

**DOI:** 10.1101/2022.11.08.515627

**Authors:** Keiji Ota, Lucie Charles, Patrick Haggard

## Abstract

Humans and some other animals can autonomously generate action choices that contribute to solving complex problems. However, experimental investigations of the cognitive bases of human autonomy are challenging, because experimental paradigms typically constrain behaviour using controlled contexts, and elicit behaviour by external triggers. In contrast, autonomy and freedom imply unconstrained behaviour initiated by endogenous triggers. Here we propose a new theoretical construct of adaptive autonomy, meaning the capacity to make behavioural choices that are free from constraints of both immediate external triggers and of routine response patterns, but nevertheless show appropriate coordination with the environment. Participants (N = 152) played a competitive game in which they had to choose the right time to act, in the face of an opponent who punished (in separate blocks) either choice biases (such as always responding early), sequential patterns of action timing across trials (such as early, late, early, late…), or predictable action-outcome dependence (such as win-stay, lose-shift). Adaptive autonomy was quantified as the ability to maintain performance when each of these influences on action selection was punished. We found that participants could become free from habitual choices regarding when to act and could also become free from sequential action patterns. However, they were not able to free themselves from influences of action-outcome dependence, even when these resulted in poor performance. These results point to a new concept of autonomous behaviour as flexible adaptation of voluntary action choices in a way that avoids stereotypy. In a sequential analysis, we also demonstrated that participants increased their reliance on belief learning in which they attempt to understand the competitor’s beliefs and intentions, when transition bias and reinforcement bias were punished. Taken together, our study points to a cognitive mechanism of adaptive autonomy in which competitive interactions with other agents could promote both social cognition and volition in the form of non-stereotyped action choices.

## Introduction

The capacity for voluntary action is often considered a defining feature of the human mind, but what volition is remains controversial. Two strong aspects of the definition of volition rely on exclusion: actions that are immediate responses to an external triggering stimulus are less voluntary than actions that are internally generated; actions that merely continue a routine, stereotyped generative pattern are less voluntary than actions that are planned anew, or actions that arise spontaneously. Both of these key features of volition are lacking in laboratory studies of behaviour, which traditionally seek tight experimental control through imperative stimuli and overlearned response patterns.

Many studies of volition involve a paradoxical instruction to act in a way that is endogenous and spontaneous (Brass & Haggard, 2007; Fleming et al., 2009; Jahanshahi et al., 1995; Libet et al., 1983) without being triggered by external factors (Jenkins et al., 2000). This minimalist approach can elicit actions that are stimulus-independent. Often, it leads to actions that have an unpredictable or random pattern, either because this is explicitly instructed (Baddeley et al., 1998; Baddeley, 1966; Jahanshahi et al., 2000), or because the participant implicitly intuits that stereotypy and voluntariness are opposites. But such studies neglect the reasons or goals that an agent may have for acting, and omit the crucial ‘Why?’ aspect of voluntary action (Haggard, 2019; Shadlen & Roskies, 2012).

Philosophical accounts of free action distinguish between freedom **from** external constraint, and freedom **to** act in accordance with goals and desires (Berlin, 1969; Bonicalzi & Haggard, 2019). Here we propose that competitive game scenarios can capture both the “freedom from” aspect of voluntary action (since the game structure can require the participant to act in the absence of triggering stimuli, or stereotyped response patterns) and “freedom to” aspect (since the participant has the clear goal of defeating the opponent and achieving good performance). For example, in the ‘rock, paper, scissors’ game, each participant selects an action without first seeing the action of their opponent. Second, many competitive games encourage unpredictable action over stereotyped action, even without any explicit instruction to act randomly. In addition, many games have a “means-ends” structure, where active selection between alternative possible action choices is the key to success. In order to defeat the opponent, each participant needs to avoid stereotyped exploitative behaviours and instead explore alternative strategies. Humans and non-human primates indeed respond to competitive pressure by initiating exploratory behaviour (Barraclough et al., 2004; Lee et al., 2004; Lee et al., 2005).

In the present study, we have studied a classical paradigm of volition through the lens of a competitive game. Participants chose when to make a simple manual keypress action (Libet et al., 1983), while trying to avoid colliding with a competitor. We designed several virtual competitor algorithms, each designed to predict and punish stereotyped choice patterns associated with a particular kind of cognitive strategy for action generation, pressurising players to become free from the corresponding pattern. The ability to adapt voluntary action choices under various competitive pressures offers quantitative measures of individual autonomy. These measures can identify how free the agent can become from specific action-generation strategies or cognitive habits and how successful they are in remaining free to achieve their goal (of succeeding in the game). For example, if people are unable to avoid repeating the same action choice, even when the current game environment punishes them for doing so and thus provides good reasons to innovate actions, one might question in what sense their actions are truly free. This conceptualization of free and autonomous action is widely used in understanding addiction (Weinberg, 2020; Weinberg, 2022).

We conceptualised three distinct expressions of autonomous behaviour, corresponding to three different constraints from which their action should become free. First, people may have a bias towards simple repetition of a given action choice (Dolan & Dayan, 2013; Robbins & Costa, 2017). We refer to this pattern of behaviour as *choice bias*. Consider the simple task of repeatedly generating one of three digits (see Figure 1A). Agent X may prefer to choose “1”, for whatever reason. Suppose now that a competitor punishes X for repeating one choice within a game scenario. If X’s action is reasons-responsive to the environmental challenge, agent X should now choose the two other digits more often and should approach an agent who has no preference for any options. This adaptive capacity may reflect the autonomy each agent has over their choice bias. We call this quantity *adaptive autonomy*.

**Figure 1.**
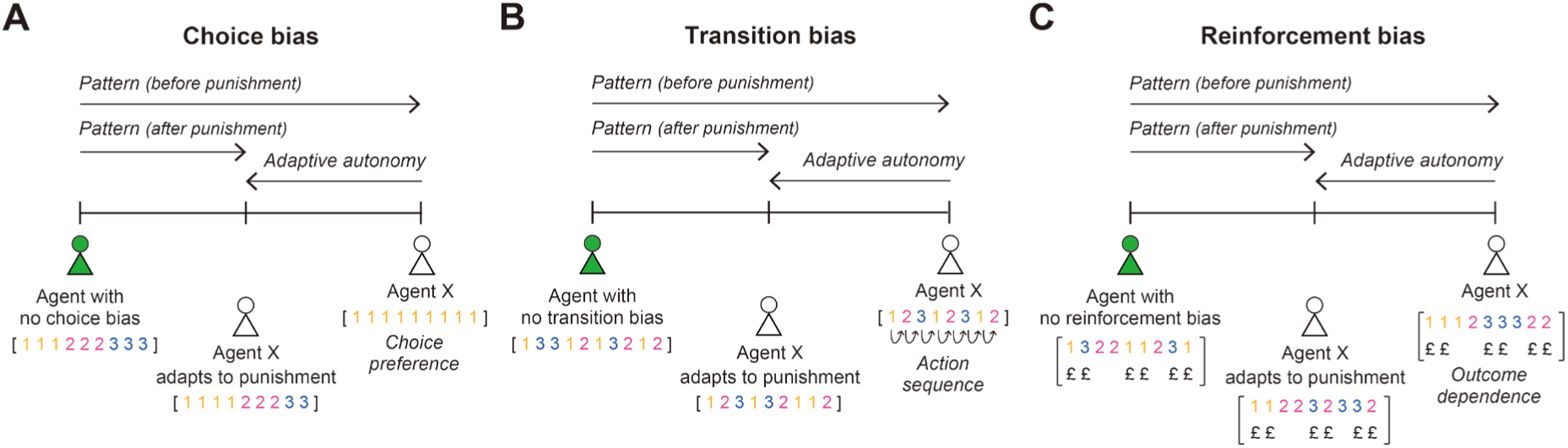
Hypothetical experiment. An agent is required to generate digits 1, 2, 3, as shown in the square brackets. **A**. Agent X exclusively selects the digit 1. They have a *choice bias*. In contrast, an agent with no choice bias selects each digit equally often. **B.** Agent X generates successive digits by counting. In contrast, an agent with no transition bias does not use any obvious transition strategy between moves. **C.** Agent X tends to repeat any action that has just been rewarded. In contrast, an agent with no reinforcement bias does not use information about the outcome of the present action to decide the next action. In each case, the leftward arrow indicates the reduction in a given bias when the opponent punishes that pattern of action choice by using it to predict the agent’s next move, i.e., adaptive autonomy.

In the second environment, the competitor pressurised participants to change any habitual action chains or routines (Lashley, 1951; Robbins & Costa, 2017; Rosenbaum et al., 2007). In our example, integer counting (“1, 2, 3”) is such a choice pattern (agent X in Figure 1B). We refer to this as *transition bias*. The only way to completely avoid such biases is to generate each choice independently from the previous trial. Yet studies of random number generation show people find this difficult (Baddeley et al., 1998; Ginsburg & Karpiuk, 1994). Figure 1B illustrates potential behavioural adaptation to punishing transition bias.

Lastly, we consider how people respond to previous action successes and failures, by considering *reinforcement bias*. In Figure 1C, agent X repeats the digit that has been rewarded. In contrast, we can imagine an agent who generates each choice independently from whether the previous outcome was rewarded or not. Most reinforcement learning approaches assume that a ‘win-stay lose-shift’ strategy is natural, or even unavoidable (Worthy et al., 2013). Here we test whether people can unlearn this familiar pattern of outcome-dependent action choices when it is punished by a competitor. Adaptive autonomy would mean that an agent would be able to break the association between the previous outcome and the forthcoming action (Fig. 1C). Such freedom from influences of past reinforcement is considered important in situations such as combatting addiction, and recovery from trauma.

In addition, we considered whether adaptive autonomy with respect to these three different forms of constraint could be controlled by a shared cognitive mechanism (Braver, 2012; Braver et al., 2007; Tang et al., 2022), or whether there are distinct cognitive control modules specific to modulating each particular bias. To do this, we explored correlations across individuals among the three adaptive autonomy measures derived from these three punishment schedules.

Finally, we modelled the learning process by which people generated a new action in order to avoid colliding with the competitor. Due to the nature of the competitive game, the participant could potentially develop a model of the opponent by understanding what options the opponent is going to select, and thus improve their beliefs about the opponent’s behaviour and guide their own action choices accordingly. We thus examined whether participants moved from model-free reinforcement learning (Dolan & Dayan, 2013) towards model-based behaviour. This would represent an interesting convergence between theory of mind, voluntary action, and behavioural autonomy.

## Methods

The full description of the data analysis and computational models can be seen in Supplementary Material.

### Participants

One hundred and fifty-nine participants (age range = 18–45, M = 29.5 yo, SD = 7.2) were recruited online via the Prolific website (https://www.prolific.co/). Participants received a basic payment of £3.75 for their participation in a 30-minute experiment. They earned a bonus of up to £4 based on their performance on the task. There were 95 female participants and 64 male participants. Recruitment was restricted to the United Kingdom. Seven participants were excluded from the analysis due to insufficient performance and the remaining 152 participants were analysed. All procedures were approved by the Research Ethics Committee of University College London. Participants gave informed consent by checking and validating the consent form.

### Experimental design

#### Apparatus

We used the JavaScript library jsPsych (de Leeuw, 2015) and the plugin jsPsych-psychophysics (Kuroki, 2021) to program the task and hosted the experiment on the online research platform Gorilla (https://gorilla.sc/) (Anwyl-Irvine et al., 2020), which participants could access through their browser on their own computer. We assumed that monitor sampling rates were typically around 60 Hz, with little variation across computers (Anwyl-Irvine et al., 2020). The size and position of stimuli were scaled based on each participant’s screen size which was automatically detected. The size of stimuli reported below are for a monitor size of 15.6” (view point size, width x height: 1536 x 746 pixels). A short version of our task is available to play online: https://research.sc/participant/login/dynamic/5D39406C-8AB9-4F6D-8FD7-9F995D9DCB83

#### Stimuli and task

Each trial started with a fixation cross, which appeared for 0.6–0.8 seconds. The images of a tree, a flock of birds and a basket containing apples then appeared (Fig. 2A). A tree (width x height: 307 x 375 pixels) was shown on the left of the screen and a flock of birds (width x height: 30 x 22 pixels each) were located on the tree. A rectangular basket of apples (width x height: 153 x 167 pixels) was presented in the bottom centre. After the fixation cross disappeared and all images appeared, the participants were given 4.5 sec to throw the food. Pressing a key initiated delivery of the food to a storage location which was located at 447 pixels forward from the start point. This delivery took 1.5 sec. We programmed the birds to attempt to intercept and catch the food. The birds on each trial were designed to intercept the food thrown within one of three intervals: 1) early throw (0–1.5 sec), 2) middle throw (1.5–3.0 sec) or 3) late throw (3.0–4.5 sec). After their departure, it took approximately 0.25 sec for each bird to reach and pass through the storage location. The participants competed with the virtual competitor, aiming to deliver food before or after the birds reached the storage location. We counted whether one of the birds overlapped with the food when the delivery was completed. If this was the case, the food was caught, and the participant lost on that trial. If not, the food was delivered without it being caught, and the participant won the trial. If no response was submitted before 4.5 sec, the food was launched automatically, and a trial was terminated as a timeout. Finally, we provided a feedback message: “Success!”, “Fail!” or “Timeout!”, which lasted for 1.0 sec. The next trial then started with a fixation cross.

**Figure 2.**
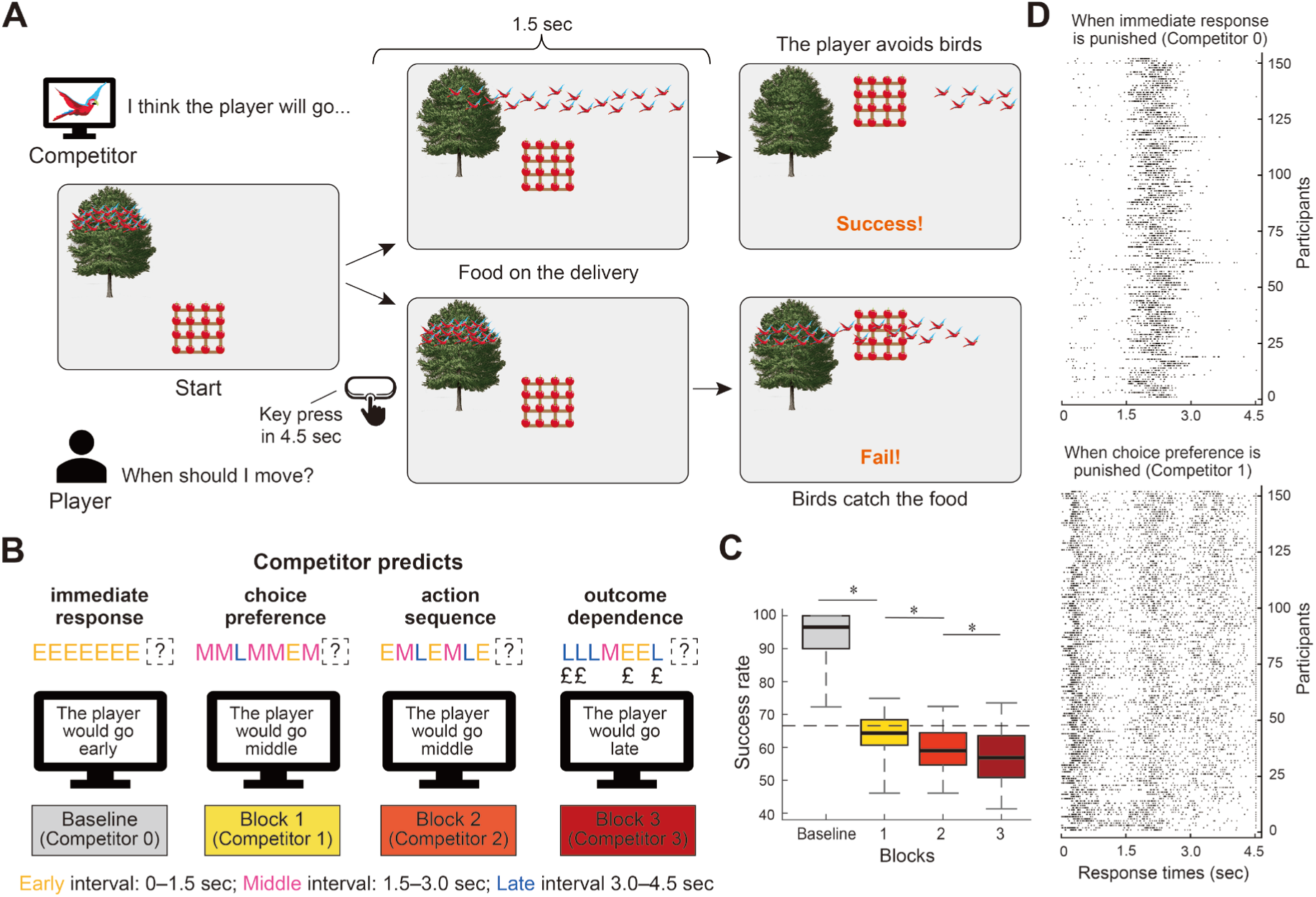
Virtual competitive environments. **A**. A trial sequence. A participant decides when to throw food within a 4.5 second time window. The food is delivered at the top-centre of the screen 1.5 sec after their key press. A virtual competitor (i.e., a flock of birds) attempts to intercept the food by adjusting the time at which it flies out of a tree. Participants win a trial if they throw at a time which avoids the food being caught by the birds. **B**. Experimental design. On each trial, the virtual competitor intercepted the food during one of three action intervals, 1) early throw (0–1.5 sec), middle throw (1.5–3.0 sec) or late throw (3.0–4.5 sec). In the baseline block, the early interval, associated with immediate responding, was punished. In block 1, *choice bias* that favoured any one interval over all others was predicted and punished. In block 2, *transition bias* (i.e., any sequential pattern) was punished. In block 3, *reinforcement bias* (i.e., any outcome dependence) was punished. **C**. Success rate of avoiding the birds against each type of competitor. A dashed line denotes the chance level. For each box, the central mark represents the median, the edges of the box are the 25th and 75th percentiles and the whiskers are the 2.5th and 97.5th percentiles. * *p* < significant level after Bonferroni correction, Wilcoxon signed rank. N = 152. **D**. Response times before the punishment of choice bias in the baseline block (upper panel) and during the punishment in block 1 (lower panel). The response time data for each participant are aligned in each row. Each small dot represents a reaction time in each trial.

In the instructions, we emphasised the following points. First, merely reacting to the moment when a stimulus is absent – the birds resting in the tree – will not win the game because the birds can travel much faster than the food. Second, merely waiting for the birds to pass is not a solution because of the time constraint. Third, the birds’ flying interval is not the same on every trial, nor is it random. Instead, the birds can learn when the participant tends to act. Therefore, predicting when the birds will likely fly out of the tree and randomising the time to throw is important. The participants were not told explicitly about the strategy used to decide the bird’s time of flight.

#### Procedure

Participants first received the instructions and viewed a set of demonstrations about the task. Following some practice trials, the participants completed four blocks of the game with a 1-minute break between blocks. The baseline block lasted 2.5 minutes while the remaining blocks 1, 2 and 3 lasted 5 minutes each. The participants got as many throws of the food as they could in the 2.5 or 5 minutes. The participants could check remaining time in each block. We used time, and not trial number, to terminate each block so that participants did not respond immediately on every trial, finishing the game early. The bonus payment was determined by the percentage of throws that successfully avoided birds and was paid up to £1 for each block: if 40 out of 60 throws were successful, we paid £1 x 66.6% = £0.66 (average bonus, baseline: £0.94; block 1: £0.63; block 2: £0.59; block 3: £0.57). The success rate and the timeout rate were included in the feedback. Nevertheless, we assumed that some participants might consume time by not focusing on the game. To prevent this, we encouraged participants to sustain the proportion of timeout trials under 5%.

### Competitor design

The virtual competitor design was primarily inspired by studies with primates (Barraclough et al., 2004; Lee et al., 2004) and rodents (Tervo et al., 2014). We programmed the learning algorithm (i.e., birds) to seek out behavioural patterns in the participant’s choice history and to pressure participants into novel behaviour. The participants could decide the time to act between 0 sec (as soon as birds and food appeared) and 4.5 sec (until timeout). To make the competitor’s prediction simple, we discretised the time window into three intervals, 1) early interval (0–1.5 sec), middle interval (1.5–3.0 sec) and late interval (3.0–4.5 sec). The participants were not informed about this discretization. Given past behaviour, the competitor predicted which interval a participant was likely to select. If the participant threw the food during the interval predicted by the competitor, the participant lost. If the participant threw the food during one of two other intervals, the participant won.

We designed four distinct competitors (Fig. 2B). First, in the baseline block, Competitor 0 punished participants for responding immediately. In this block, the birds simply blocked the early throw on every trial. Thus, the stimulus-absence behaviour corresponded with waiting until the middle interval. Competitor 0 thus pressured participants to use voluntary control to resist immediate response (Haggard, 2019).

Second, in block 1, Competitor 1 made predictions based on the participant’s currently-preferred interval and punished *choice bias*. On each trial, a history of the participant’s past ten choices was used to estimate the probabilities of selecting the early, middle and late interval. The choice probabilities were then used to generate the competitor’s prediction for the upcoming choice. For instance, if the participant chose the early interval seven times, the middle interval twice, and the late interval once, the competitor penalised the early interval 70% of the time, the middle 20% of the time and the late 10% of the time. Thus, Competitor 1 required participants to distribute their choices across time.

In block 2, Competitor 2 sought out *transition bias* or sequential pattern between the choice of interval on one trial and the next. A history of the past 60 trials was used to estimate the conditional probabilities of selecting three intervals given the previous choice. Competitor 2 then exploited these conditional probabilities to predict the upcoming choice given the last choice. Competitor 2 pressured participants into avoiding habitual transition patterns.

Finally, in block 3, Competitor 3 punished *reinforcement bias* (outcome dependence) – which interval the participant is going to select after the participant made a particular response and won a trial or lost a trial –. Competitor 3 used the same search algorithm as Competitor 2 with the exception that they looked for the conditional probabilities of selecting three intervals to predict the upcoming choice given the last choice and the last outcome. Competitor 3 required participants to act independently from the previous outcome. Supplementary Material describes how we programmed different computer algorithms.

### Data analysis

Because the birds’ decisions were discretized into one of the three response intervals, we similarly discretized participants’ reactions into 1) the early response: responding in 0–1.5 sec, 2) the middle response: responding in 1.5–3.0 sec, 3) the late response: responding in 3.0–4.5 sec (including timeout). Note that these intervals were not known to the participants, who experienced a continuous temporal interval on each trial.

#### Quantifying decision bias scores

Statistical distance is a standardised way of measuring the extent to which the observed probability distribution is different from the target probability distribution. We calculated the Kullback-Leibler divergence to quantify the extent to which the participant’s choice probability distribution is different from the choice probability distribution that a bias-free agent would exhibit. See Figure 3.

**Figure 3.**
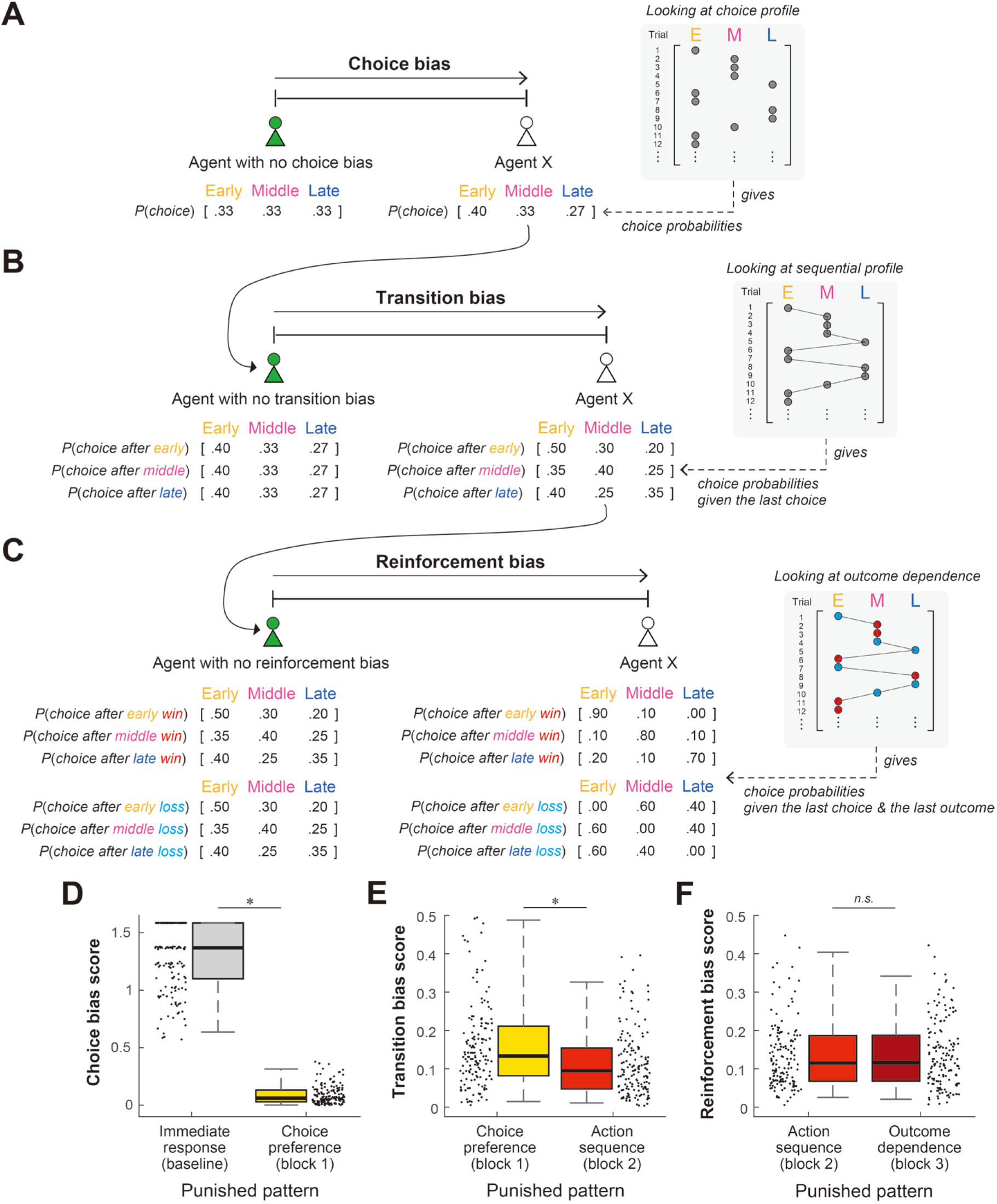
Measuring decision bias scores with respect to different punishments. In the raster plot, potential actions for the early, middle and late intervals are shown. A vector of the numerical values illustrates the probabilities for choosing these three actions. **A.** Agent X selects the early interval more frequently than the other two intervals. In contrast, an agent without choice bias selects each interval equally often. We measured the statistical distance between individual and a bias-free agent with respect to a pattern similarity between their choice probabilities. **B.** The dashed line from the choice bias vector in A to the transition vector in B shows the logical relation between choice and transition: any change in choice bias inevitably changes the transition vector. Transition bias is considered as the additional change over and above such basic effects of the choice bias. An agent without any transition bias should determine their next action independently from the previous action. **C.** The dashed line from the transition bias vector in B to the reinforcement vector in C shows the logical relation between transitions and reinforcements. Reinforcement bias is considered as the additional change over and above such basic effects of transition bias. An agent without any reinforcement bias should determine their next action independently from the previous success or failure. **D-E.** Planned comparisons show the degree of adaptive autonomy (bias reduction) when choice bias is punished (D), when transition bias is punished (E) and when reinforcement bias is punished (F). * *p* < significant level after Bonferroni correction, Wilcoxon signed rank. On each box, the central mark represents the median, the edges of the box are the 25th and 75th percentiles and the whiskers are the 2.5th and 97.5th percentiles. Dots adjacent to each box show individual scores.

1) Choice bias. The probabilities of choosing the early, middle and late interval for a bias-free agent would be 0.33, respectively. We computed the choice probabilities *P*(*c*) given a history of intervals each participant chose in each block. The K-L divergence is then

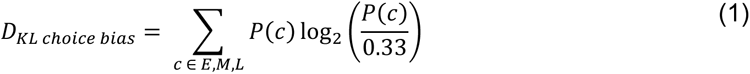

2) Transition bias. We computed the conditional probabilities of choosing the early, middle and late interval given the interval chosen on the previous trial *P*(*c|c*_−1_). We measured the K-L divergence of these participant’s conditional probabilities from the participant’s choice probabilities. The K-L divergence for each previous interval *c*_−1_ is computed as

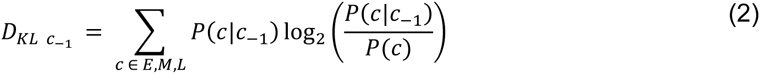

The total K-L divergence as a weighted sum is then

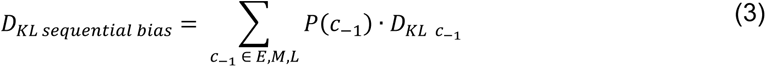

We quantified the deviation of patterns associated with the previous choice from sequential patterns logically expected from the participant’s own choice probabilities. Competitor 2 specifically detected and punished this conditional dependence, on the top of choice bias in which Competitor 1 punished.

3) Reinforcement bias. We computed the conditional probabilities of choosing the early, middle and late interval given the interval chosen and the outcome obtained on the previous trial *P*(*c|c*_−1,_ *o*_−1,_). We measured the K-L divergence of these participant’s conditional probabilities from the participant’s conditional probabilities given the previous interval solely. The K-L divergence for each previous interval *c*_−1_ and each previous outcome *o*_−1_is computed as

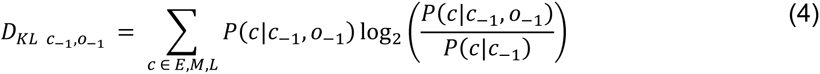

The total K-L divergence as a weighted sum is then

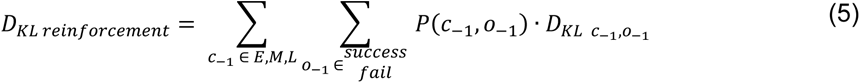

This way, we quantified the deviation of patterns associated with both the previous choice and the previous outcome from patterns logically expected from the conditional dependence on the previous choice alone. Competitor 3 specifically detected and punished this outcome dependence, in addition to the choice bias and transition bias punished by Competitor 2.

#### Statistical analysis

We tested the performance difference by Wilcoxon signed rank test. The alpha level of 0.05 was corrected by the number of tests we performed in each class of test (Bonferroni correction).

### Lagged correlation

We performed an ordinary regression analysis to examine the extent to which the participants’ action choices (early, middle or late response) were influenced by their own previous actions and their opponent’s previous actions (Devaine et al., 2014; Devaine et al., 2017). We fitted a cumulative link mixed-effects model with the current choice *c* as the dependent variable. The independent variables were one’s own previous choices *c*_−1_,…, *c*_−5_ up to the 5th preceding trial, the opponent’s previous choices *c′*_−1_,…,*c′*_−5_ up to the 5th preceding trial and a dummy variable which indicates block 1 (choice block), block 2 (transition block) or block 3 (reinforcement block). Performance on the baseline block was removed from the analysis since choice transitions between trials were rare, as participants generally resisted immediate responses. A dummy variable interacted with each lagged choice to test whether these lagged correlations changed across blocks 1, 2 and 3. Slopes were allowed to vary between participants as random effects. A mixed-effect model was fit using *ordinal* package (Christensen, 2022). In addition, an ordinary regression was performed on the sequence of action choices simulated by computational models (see *Computational models* in

Supplementary Material).

## Results

### Experimental task

Participants were asked to decide when to press a key that caused some food to be delivered to a storage location. They were competing with a virtual competitor, represented as a flock of birds (Fig. 2A). The birds tried to intercept and catch the food during the delivery process, by deciding when to fly out of a tree. The participant’s task was to deliver the food without it being caught by the birds. We programmed the birds to predict the time of the participant’s next action based on the history of their reaction times. Based on this prediction, the birds made a choice of when to fly on each trial. They flew at a time that was designed to intercept the food thrown by a participant within one of three intervals: 1) early throw (0–1.5 sec), 2) middle throw (1.5–3.0 sec) or 3) late throw (3.0–4.5 sec). Participants were not told whether the birds chose a continuous or a 3-option distribution. The participants could win a trial by pressing the key during one of two intervals that the birds did not select. The intervals were not explicitly demarcated for the participant, who experienced a continuum of potential action times in each trial. Participants could not perform the task reactively because the birds could travel much faster than the food. If the participants simply waited for a moment when no birds were present and then threw in reaction to that moment, the birds could suddenly appear and intercept the food. Therefore, the participants were instructed to predict when the birds would appear and to try to avoid them. This feature means that our participants’ actions were stimulus-independent.

There were 4 blocks in total. In each block, the participants competed with a class of competitors that pressurised a distinct constraint or a stereotypical action-generation strategy (Fig. 2B). In the baseline block, Competitor 0 was programmed to punish the participants for responding immediately: the birds consistently punished a participant who threw in the early interval, so the participant was incentivized to wait to avoid being intercepted. In block 1, Competitor 1 punished choice bias: if a participant selected one interval more often than the other two, then the birds became more likely to intercept the food during the same interval. In block 2, Competitor 2 predicted transition bias and punished any predictable association between the time of the participant’s current throw and the time of the preceding throw. Finally, in block 3, Competitor 3 punished reinforcement bias by seeking out whether the time of the current throw was associated with both the time of the preceding throw and the preceding outcome. Thus, the participants required progressive degrees of autonomy across blocks: they needed in each block to act in a way that was even more unconstrained than required by the competitors they had played previously.

Participants were never told when they should act on any given trial. Participants did not receive any explicit instruction or explanation about how the competitor would behave or respond to the participant in each particular block. Instead, they could only monitor the success/failure of avoiding the birds on each trial, and adapt their behaviour accordingly to try to avoid the birds on future trials. Thus, successful performance under different punishment regimes would depend on implicit mechanisms of behavioural adaptation rather than explicit understanding.

We first checked the percentage of successful bird-avoiding trials. The participants achieved near-perfect success rates against Competitor 0 who simply punished immediate responding (Fig. 2C; Median [Mdn] = 96.6%). The participants avoided an immediate response and initiated the throw with a mean of 2.15 sec (SD 0.49 sec) after the trial started (Fig. 2D upper panel). In block 1, the success rate dropped to 66.6%, which, interestingly, would be predicted from selecting one of the three intervals at random (Fig. 2C; Mdn = 64.3%, *p* < .001, *z* = 10.69 for blocks 0 versus 1, Wilcoxon sign rank). The success rate further decreased in block 2 (Fig. 2C; Mdn = 59.0%, *p* < .001, *z* = 5.74 for blocks 1 versus 2, Wilcoxon sign rank) and even further in block 3 (Fig. 2C; Mdn = 56.9%, *p* = .015, *z* = 2.44 for blocks 2 versus 3, Wilcoxon sign rank). Therefore, our progressive series of punishments increased the predictive power of punishing the participants’ responses.

### Do people adapt to punishments of habitual actions?

We next examined the extent to which people could adapt to punishment of different behavioural biases. We therefore developed a measure that reflects an individual’s tendency towards the pattern of action selection for punishment on each block. We measured the statistical distance (Kullback-Leibler [K-L] divergence) between the observed probabilities of selecting the early, middle and late intervals, and the probabilities that an agent, who does not have a particular type of bias would exhibit (Figure 3A-C). A lower statistical distance means that the individual is similar to this bias-free agent. We call this quantity a *decision bias score*. The decision bias score *before* the punishment of a specific pattern reflects an individual’s trait regarding when to act. We then looked at the *change* in decision bias scores relative to the previous block, when a given individual pattern was punished. We quantified the score for choice bias, transition bias and reinforcement bias, respectively (Figure 3A-C). A greater change in bias score would indicate stronger *adaptive autonomy*, or ability to update reasons-guided voluntary action choice when a habit becomes punishable.

Participants began to distribute action times appropriately when Competitor 1 started punishing the repetition of the same action interval (Fig. 2D). Accordingly, their choice bias score**—**a statistical distance of the observed choice probabilities from probabilities 0.33 (Fig. 3A)**—**reduced after punishment (Fig. 3D; Mdn = 1.37 for the punishment of immediate response versus Mdn = 0.06 for the punishment of choice bias, *p* < .001, *z* = 10.69, Wilcoxon sign rank). Competitor 1 did not penalize transition patterns from one action to the next and allowed participants to still use *transition bias*. We quantified the transition bias score by considering the extent to which probabilities given the previous action are explained by one’s own choice probabilities (Fig. 3B). We found that the transition bias decreased after the competitor pressurised transition patterns (Fig. 3E; Mdn = 0.13 for the punishment of choice bias versus Mdn = 0.09 for the punishment of transition bias, *p* < .001, *z* = 3.82, Wilcoxon sign rank). Against Competitor 3, the participants were asked to act even more freely to avoid patterns following positive or negative outcomes. We evaluated the reinforcement bias score by considering the extent to which probabilities given the combination of previous action and the previous outcome are explained by probabilities given the previous action alone (Fig. 3C). The reinforcement bias did not show any significant improvement under competitor punishment (Fig. 3F; Mdn = 0.11 for the punishment of transition bias versus Mdn = 0.11 for the punishment of reinforcement bias, *p* = .79, *z* = 0.27, Wilcoxon sign rank).

We further tested the possibility that the participants adapted differently to the influence of positive and negative reinforcement. The processes that follow a positive outcome may be different from those that follow a negative outcome (Gehring & Willoughby, 2002; Hajcak et al., 2006; Vickery et al., 2011), leading to a stereotypical win-stay lose-shift strategy (Wang et al., 2014). We quantified the positive reinforcement bias and the negative reinforcement bias score separately (SFig. 1A). Nevertheless, we did not find significant improvements in the positive reinforcement bias (SFig. 1B; Mdn = 0.29 for the punishment of transition bias versus Mdn = 0.34 for the punishment of reinforcement bias, *p* = .21, *z* = -1.26, Wilcoxon sign rank) nor in the negative reinforcement bias (SFig. 1B; Mdn = 0.43 for the punishment of transition bias versus Mdn = 0.41 for the punishment of reinforcement bias, *p* = .19, *z* = 1.30, Wilcoxon sign rank). These results suggest that people are able to become more autonomous from standard habitual choices and habitual action transitions but cannot break away from outcome dependencies. That is, people display stereotyped patterns of being guided by reinforcement, such as win-stay lose-shift, even when they are discouraged to do so.

### Is there a common factor underlying all adaptive autonomy?

The bias reduction between the pre-punishment and the post-punishment phases provides a measure of adaptive autonomy for each decision bias (Fig. 3D-F). Theories of domain-general cognitive control propose that proactive, strategic cognitive control acts similarly across cognitive tasks (Braver, 2012; Braver et al., 2007; Tang et al., 2022). The adaptive ability to become free of one kind of bias would be expected to correlate with the ability to become free of another bias, to the extent that both depend on a common, domain-general mechanism. In fact, the correlation structure of the changes in our decision bias scores showed poor correlations across our 152 participants (Figure 4; *r* = -0.07∼-0.01). This suggests that the ability to voluntarily regulate one bias is not strongly associated with the ability to regulate other biases. This also suggests that our measurements are separable and evaluate three distinct forms of autonomy conceptualised above, rather than a single common form of autonomy. Since this correlation approach would be undermined by a floor effect of participants who showed no adaptive autonomy at all, we repeated the correlations including only participants who showed a numerical change in decision bias. We did not find significant correlations among adaptive autonomy to the three bias classes even in this restricted group (*r* = -0.06∼0.04). We also computed Jeffreys–Zellner–Siow Bayes factor for correlation (Wetzels & Wagenmakers, 2012). The Bayes factor *BF_10_* using the JZS prior was 0.07 (choice versus transition), 0.06 (choice versus reinforcement), and 0.09 (transition versus reinforcement), respectively. Bayes factor indicates that the data strongly favours a null hypothesis of independence of variables we tested (Jeffreys, 1961). These results suggest that people recruit distinct cognitive capacities to exert autonomous behaviour when unlearning choice bias, transition bias and reinforcement bias, contrary to concepts of a general cognitive control capacity.

**Figure 4.**
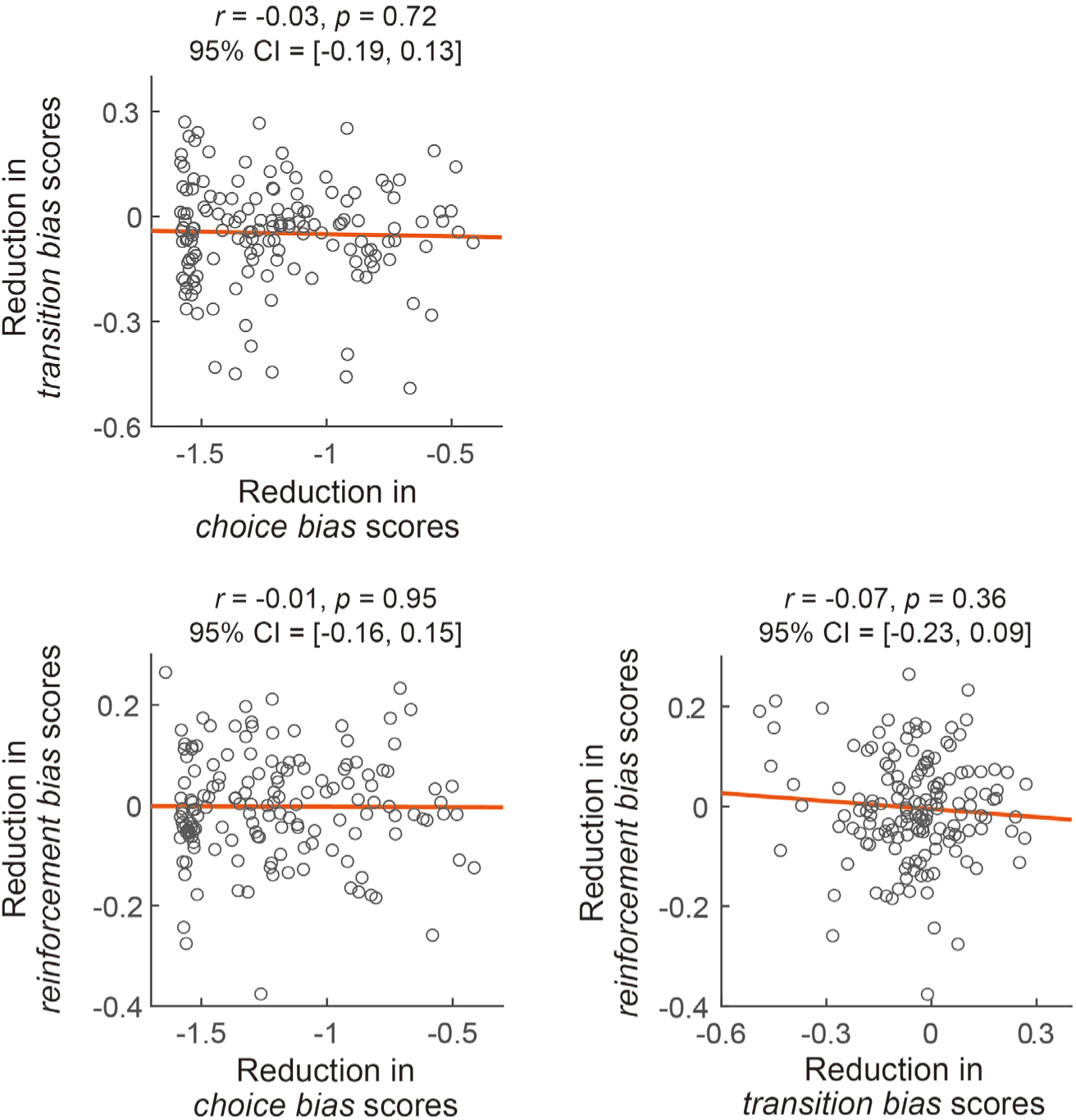
Correlation structure underlying measures of adaptive autonomy. A planned comparison between the pre-punishment and the post-punishment phase in Figure 3D-F provides bias reduction (i.e., adaptive autonomy) for each class. Across 152 participants, there is no strong correlation between the three measures of adaptive autonomy.

### A mechanism that underlies adaptive autonomy

We demonstrated that people become less stereotypical and more volitional by adapting to external constraints of competitive pressures. In competitive and strategic settings, voluntary actions that aim a goal-directed and less stereotyped behaviour can be achieved via two dissociable ways. The first is a stochastic approach which aims to increase the entropy of behaviour. This approach does not involve any experience-based updates on action choices. A second social-cognitive approach exploits belief-learning processes (Camerer, 2003; Hampton et al., 2008; Zhu et al., 2012). Here the agent forms an explicit model of the structure of the game by learning the competitor’s likely action (1st order belief) or by learning how the competitor predicts the agent’s likely action (2nd order belief). Broadly speaking, the algorithm adopted by the virtual competitor in this task is considered 1st order belief learning. By knowing that the competitor possesses the 1st order belief and predicts how the agent themselves will act, the 2nd order belief learner generates an action that breaks the competitor’s prediction.

Both the stochastic account and socio-cognitive account could potentially form part of a theory of volition since they both involve a choice between alternative possible actions, and both are goal-directed. When human agents try to produce statistical randomness, they rely heavily on working memory resources (Baddeley et al., 1998). The socio-cognitive approach involves learning and exploiting a model of the structure of the game in order to select the best action at a particular trial. On one view, both first-order and second-order belief learning involve the element of mentalising because they simulate the competitor’s intention (Hampton et al., 2008; Zhu et al., 2012). On another view, first-order belief learning is interpreted as a special case of reinforcement learning which does not involve learning about the competitor’s mental state (Abe & Lee, 2011; Camerer & Ho, 1999) since the underlying mathematical formula is equivalent to adjusting action values for both chosen and unchosen actions according to both their actual rewards and hypothetical, fictive rewards that the agent would have observed. Nevertheless, learning successful actions from fictive rewards still involves the element of learning the overall reward structure of the game.

To test whether our participants relied more on purely stochastic processes or socio-cognitive processes, we examined patterns of sequential dependence of the data (Devaine et al., 2014; Devaine et al., 2017). Specifically, we simulated a sequence of choices generated by the agent who plays against the competitor (see *Computational models* in Supplementary Material). We then regressed their simulated choices onto their own past choices and their opponent’s past choices. As illustrated in Figure 5A, an agent who uses a purely stochastic process has no sequential structure to their action choices and their opponent’s choices. An agent who relies on reinforcement learning (Sutton & Barto, 2018) will tend to repeat their past choices and therefore show a sequential structure characterised by positive lag correlations with their own actions. In contrast, an agent who exploits first-order belief learning will show negative lag correlations with their opponent’s actions, because they learn the opponent’s intentions and avoid taking the same action that the opponent showed in the past. Importantly however, such an agent does not show positive lag correlations with their own actions as strongly as reinforcement learning does because first-order belief learning does not just repeat their past choices that avoided the opponent’s action but also selects the other action that the opponent did not show. Finally, an agent who relies on second-order belief learning will not show negative lag correlations with their opponent’s actions, but will show negative lagged correlations with their own actions. This is because they know that the competitor will predict their own intentions – so they should use this knowledge to avoid repetitive action patterns that the opponent might potentially use to predict future behaviours. Therefore, the crucial marker of second order belief learning is a stronger negative lagged correlation with one’s own action.

**Figure 5.**
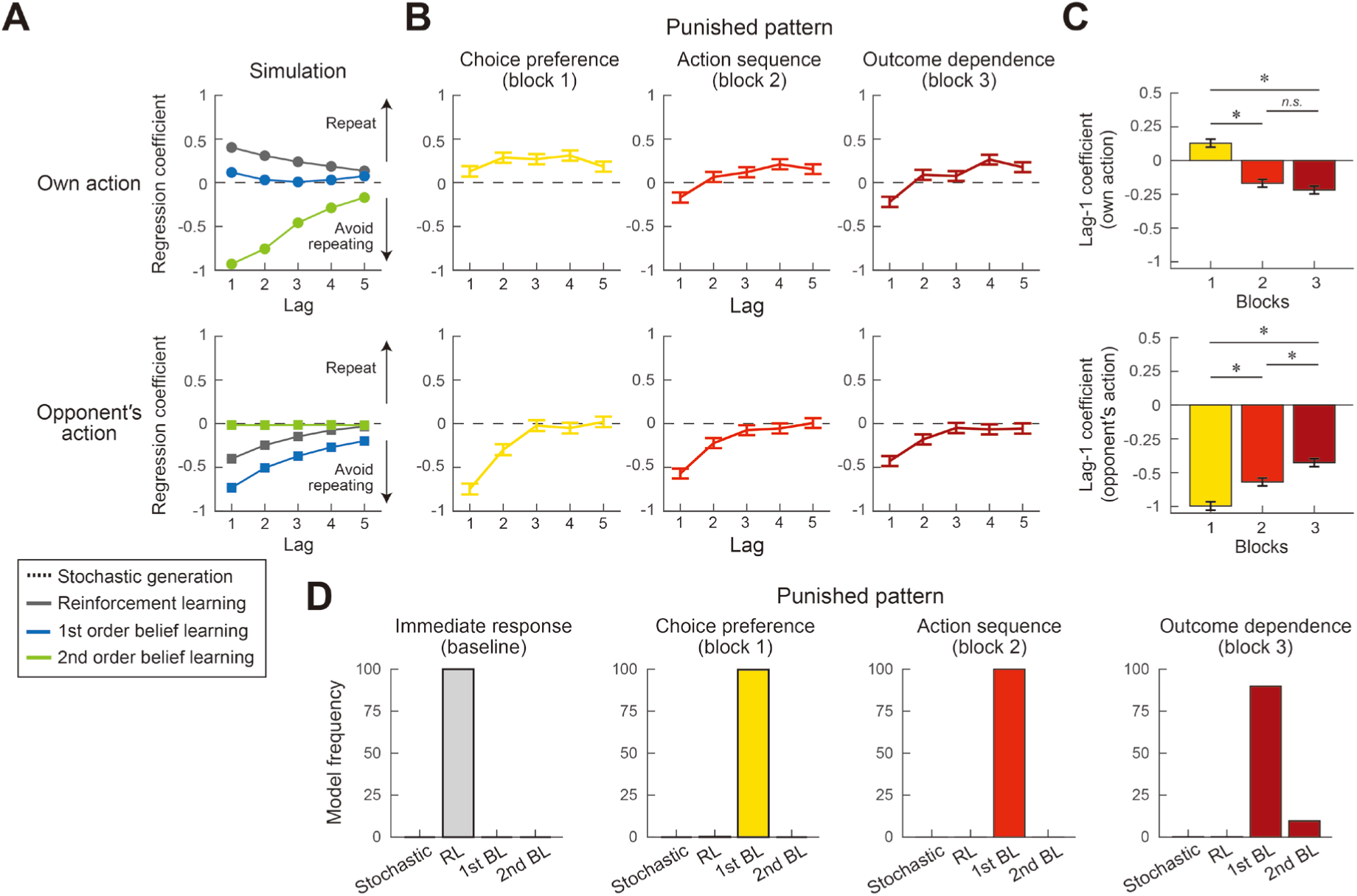
Influence of previous action choices on the current action choices. **A.** Estimated lagged correlations with the agent’s own action (upper panel) and with their opponent’s action (lower panel). In the simulation, four types of agents competed against Competitor 1. They used either stochastic action generation, reinforcement learning, first-order belief learning or second-order belief learning. The stochastic action generation process predicts no sequential dependence on both one’s own action and their opponent’s action. A positive lagged correlation with own action indicates that agents repeat the same action that agents themselves took in the past while a negative lagged correlation indicates that agents avoid repeating the same action agents took. A negative lagged correlation with the opponent’s action indicates that agents avoid taking the same action that the opponent showed in the past. **B.** Lagged correlations in participants’ choice across blocks. **C.** The values of lag-1 correlation in three different blocks are taken from panel B and replotted as a bar graph. As competitive demands increase, one’s own action in the preceding trial becomes negatively correlated with the current action (i.e., increased reliance on second order belief). * *p* < significant level after Tukey’s correction. **B&C.** Error bars represent two standard error of the mean. **D.** Estimated protected exceedance probability associated with each model. This index corresponds to the posterior probability that a given model is the most frequent one among the models we tested.

Figure 5B illustrates the lagged correlations for our participants’ data. In general, the participants’ current action was strongly correlated with the most recent action, and correlations became weaker as the lag increased. Figure 5C shows a lag-1 correlation across blocks. Participants’ own choice in the preceding trial was positively correlated with their current choice when choice bias was punished (block 1; *β* = 0.13, 95% confidence interval = [0.07, 0.19]). This pattern of result is consistent with reliance on reinforcement learning or on first-order belief learning. In contrast, this correlation was negative when transition bias was punished (block 2; *β* = -0.17, 95% CI = [-0.23, -0.11]) and also when reinforcement bias was punished (block 3; *β* = -0.22, 95% CI = [-0.27, -0.16]). Pairwise testing showed that Lag-1 correlations in blocks 2 and 3 were significantly lower than in block 1 (*p* < 0.001, *z* = 7.14; *p* < 0.001, *z* = 8.34, respectively), while there was no difference between blocks 2 and 3 (*p* < 0.44, *z* = 1.22). This pattern of results is consistent with increased reliance on second-order belief when the competitor punished transition and reinforcement biases. In addition, the opponent’s choice in the preceding trial was negatively correlated with the participant’s current action choice in all three blocks (*β* = -0.75, 95% CI = [-0.81, -0.69] for block 1; *β* = -0.57, 95% CI = [-0.63, -0.51] for block 2; *β* = -0.43, 95% CI = [-0.48, -0.37] for block 3). Pairwise comparisons showed that the negative lag-1 correlation became weaker from block 1 to block 2 (*p* < 0.001, *z* = -4.22) and again from block 2 to block 3 (*p* < 0.01, *z* = -3.55). This pattern of results is also consistent with a shift from first-order belief learning to second-order belief learning across blocks, with participants becoming progressively less dependent on the action that their opponent took on the preceding trial.

To quantify which model (stochastic action generation model; reinforcement learning, RL; 1st order belief learning, 1^st^ order BL; 2nd order belief learning, 2nd order BL) supports our participants’ behaviour, we fitted each model to predict choice sequences from the reward and from the opponent’s action choice on the preceding trial (see *Computational models* in Supplementary Material). We performed a group-level Bayesian model selection (Rigoux et al., 2014; Stephan et al., 2009) and computed the protected exceedance probability which is an omnibus measure of the probability that a given model is the most frequent one among tested models (Fig. 5D). The protected exceedance probability for the RL model to outperform the other three models was close to 100% under the punishment of immediate response at baseline. When choice bias and transition bias were punished, the estimated model frequency distribution changed and the probability for the 1st order BL model to be the most frequent model was close to 100%. When reinforcement bias was punished, the model evidence supported 2nd order BL by 9.6% while it supported 1st order BL by 90.0%. A shift from 1st-order BL to 2nd-order BL in block 3 is consistent with an increased negative lagged correlation with one’s own action and a weakened negative lagged correlation with the opponent’s action (Fig. 5 B&C).

Taken together, this suggests that the mechanism of adaptive autonomy is not simply explained by a stochastic action generation process. People do not simply express voluntary choice in competitive games by behaving more randomly. Instead, our analyses support a socio-cognitive approach which learns the first-order belief about the intention of the opponent’s action or the second-order belief about the opponent’s belief of the participant’s action. These belief learning strategies employ a model of the competitive game structure. Particularly, as the competitor’s prediction became more sophisticated, the participants’ choices increasingly relied on understanding how the competitor thinks the participant will act. This finding suggests that a cognitive mechanism of adaptive autonomy links to social cognition on the condition where active interaction with a circumstance is designed to evoke autonomous behaviour.

## Discussion

It is widely assumed that humans choose and control their own actions for reasons that are important to them. Psychologists and philosophers have argued that volition is a cluster concept which aggregates different aspects of human cognition (Haggard, 2019). However, most experimental psychology studies struggled to isolate key features of human voluntariness in the laboratory, because they used minimalist approaches which instructs participants to choose voluntarily among meaningless options. Previous neuroscientific literature has emphasized that a volitional action must be endogenous and spontaneous, rather than stereotyped. At the same time, previous philosophical literature has emphasized that a volitional action is made for a reason (Berlin, 1969). Human experimental studies have generally struggled to combine these two seemingly contradictory features of volition (Maoz & Sinnott-Armstrong, 2022; Roskies, 2010).

Here we have developed a family of novel competitive games which allow participants to act in a way which is simultaneously endogenous, non-stereotyped, and reasons-responsive. In the game, participants had to endogenously generate a new pattern of endogenous behaviour (without any immediate stimulus triggers) to thwart a competitor who aimed to punish specific behavioural patterns in each block. In this sense, our participants were reasons-responsive to the challenge represented by the current gaming environment. Voluntary actions elicited in this paradigm are taken to reflect how flexible or how adaptable reasons-guided voluntary action choices can be. By developing a series of environmental challenges and an analysis pipeline for quantifying adaptive autonomy for the first time, we found that people can become autonomous regarding both choice bias and transition bias in transition from one action to the next. However, participants had very limited ability to become free of reinforcement bias. We further showed, in a large sample, that the correlations between these three forms of adaptive autonomy were minimal. This pattern of results suggests distinct cognitive modules for these three forms of autonomy, rather than a common module or a single form. This finding contrasts with the classical vIew of cognitive control as a unitary cognitive resource that underpins willed action (Botvinick et al., 2001).

Our paradigm does not address all the aspects that might constitute volition. For instance, volition is often thought to involve consciousness. We do not know whether our participants were aware of the different competitive pressures, nor whether they strategically made less stereotypical choices because they became aware of the competitor’s rules. Future research would clarify whether adaptive autonomy relies on explicit processes involving conscious awareness of reasons for action, or on implicit processes operating outside of awareness of reasons for action.

### Measures of autonomy

Traditional experimental psychology struggles to investigate volition and autonomy, even though these are distinctive features of the healthy adult mind (Locke, 1690/1975). Experimental approaches to the study of volition often involved giving people a paradoxical instruction regarding how to behave autonomously (Baddeley, 1966; Brass & Haggard, 2007; Fleming et al., 2009; Jahanshahi et al., 1995; Libet et al., 1983). The few studies that have explicitly aimed to investigate human autonomous behaviour typically used competitive game contexts (Forder & Dyson, 2016; Wang et al., 2014; Wong et al., 2021), and have rarely considered subtypes of autonomy. We conceptualized three forms of autonomy, as freedom from three cognitively distinguishable types of action-generation strategy: *choice bias*, *transition bias* and *reinforcement bias*. We attempted to evoke autonomous behaviour by programming a competitor to punish any lack of each type of autonomy, testing these three biases in successive blocks Using a statistical distance measure derived from information geometry, we quantified the extent to which people could break their own stereotypical choice pattern when punished. We found no strong correlations between these three types of adaptive autonomy, making a single common underpinning cognitive control function unlikely, and supporting the idea that behavioural autonomy involves dissociable components. In tasks where participants react to external stimuli quickly, it has been suggested that a domain-general top-down control is used to solve different cognitive tasks (Braver, 2012; Braver et al., 2007; Tang et al., 2022). However, in free, stimulus-independent action, our results suggest that domain-specific top-down control independently regulates each particular form of autonomous behaviour: there are multiple ways to act freely, and it is therefore important to consider *from what* an agent aims to be free. We studied choice biases, transition biases and reinforcement biases, but other biases counteracting free action doubtless also exist. We showed that, for example, an agent who becomes increasingly free from choice bias may yet be unable to free themselves from the biasing effects of reinforcement.

### Relevance to classical neuropsychological tasks

Our study evokes behavioural phenomena that neuropsychologists have traditionally studied using arbitrary, open-choice tasks. For instance, our measure of choice bias is related to the capacity to inhibit a prepotent, impulsive action (Mischel et al., 1972). People usually place costs on waiting, preferring earlier rewards; a form of temporal discounting (Story et al., 2014). Next, the transition bias we measured reflects executive control and working memory which are often assessed using random number generation tasks (Baddeley et al., 1998; Jahanshahi et al., 2000). In these tasks, people perform poorly at randomising numbers, tending to rely on inappropriate heuristic rules, and misunderstanding the concept of randomness (Baddeley et al., 1998; Ginsburg & Karpiuk, 1994). These limitations are often attributed to limited working memory capacity. In contrast, the capacity to avoid the reinforcement bias is related to voluntary override of reward-seeking behaviour (Bechara et al., 1994; Lejuez et al., 2002) and to the balance between exploitation and exploration (Cohen et al., 2007). These classical executive functions are often seen as robust individual traits affecting any relevant task that draws on executive capacity (Neiman & Loewenstein, 2011; Ota et al., 2016; Ota et al., 2019).

Here we instead treat these biased action selections as a state function of cognitive control processes, which depend on the current environment, and which we therefore view as different expressions of *adaptive autonomy*. We developed a competitive game in which participants would need to avoid particular action preferences, action generating rules, or outcome dependences. Our results demonstrate that people can balance choice frequencies and break transitions between actions.

In contrast, we found people could not avoid reinforcement bias. Neither positive reinforcement bias nor negative reinforcement bias adapted when penalised. A stereotypical win-stay lose-shift behaviour has been shown in competitive games (Ota et al., 2020; Wang et al., 2014). In particular, people are less flexible regarding changing lose-shift behaviour than win-stay behaviour when adapting to new game rules (Forder & Dyson, 2016; Sundvall & Dyson, 2022). The experience of a negative outcome modulates the speed-accuracy trade-off on the subsequent trial (Desender et al., 2021; Dyson et al., 2018). Individual differences are also associated with these post-error reaction times. Individuals who make quicker decisions after a loss than after a win show a poorer performance than individuals who show the inverse pattern (Dyson, 2021). Therefore, overcoming impulsivity after failure may be a key aspect of volitional control for humans.

### Relevance to model-based behaviour

We conceptualised that a stochastic generative mechanism and a socio-cognitive mechanism could both support adaptive autonomy. However, our analyses of sequential dependence reveal that participants did not achieve adaptive autonomy by simply behaving randomly or stochastically. Rather, participants’ action choices were generated by updating action values throughout the game. In particular, trial-by-trial action generation patterns were consistent with first-order belief learning which involves predicting the opponent’s likely action. However, this interpretation could be challenged by the view that first order belief learning is merely a special case of reinforcement learning in which action values of unchosen options are updated by fictive rewards (Abe & Lee, 2011; Camerer & Ho, 1999). In any case, both interpretations involve learning the model of the environment, and therefore qualify as adaptive.

When transition bias and reinforcement bias were punished, participants increased their reliance on second-order belief learning. This strategy abandons model-free reinforcement learning and its classical win-stay lose-shift strategies. In general, as the competitor becomes more sophisticated, participants’ choices increasingly relied on understanding the competitor’s beliefs and intentions (Devaine et al., 2014; Yoshida et al., 2010) even though they were not explicitly informed about the competitor’s knowledge or action strategies in any block. In this sense, participants’ non-stereotyped action choice was linked to them spontaneously building a cognitive model of the structure of the current competitive game environment. Psychologists and philosophers have speculated on links between cognitive processes of volition and social interaction (Frith & Frith, 2023). Our study points to a cognitive mechanism of adaptive autonomy in which competitive interactions with other agents could promote both social cognition and volition, in the form of non-stereotyped action choices. We based this interpretation on a statistical approach inferring patterns of sequential dependence. Future experiments could be designed to directly manipulate participants’ social cognition and to look for its contribution to adaptive autonomy.

### Limitations

Our study has a number of limitations. First, one might question on theoretical grounds whether our paradigm offers a good testbed for volition. Somewhat similar competitive games have, after all, been proposed in studies of social cognition, decision-making and learning – without reference to volition. This criticism may be difficult to resist given that the precise definition of volition remains controversial. However, using a recent enumeration of various different features of volition (Haggard, 2019), we have shown that the present task elicits actions that are both stimulus-independent, and goal-directed, and therefore satisfy classical philosophical criteria for volition. Further, our task involves motivated selection of *when* to act, similar to a classical neuroscientific study of volition (Libet et al., 1983).

Next, our participants completed a structured series of games against progressively sophisticated competitors appearing in a fixed order. This design was logical given the hierarchical dependency of the different biases that we hypothesised could constrain the capacity for autonomous volitional action. For example, we reasoned that transition biases of action choice (block 2) could only be studied after biases in action choice per se (block 1) were first controlled for. However, fixed-order designs are subject to confounding effects of learning, fatigue and other time-dependent factors. For example, our participants’ inability to adapt to reinforcement biases (block 3) may be contaminated by an element of fatigue. While such confounds cannot be excluded absolutely by our design, our findings of successful adaptation initially (blocks 1 & 2) followed by less successful adaptation thereafter (block 3), rule out simple time-dependent or exposure-dependent effects such as fatigue. We addressed the concerns about fixed order design in another pre-registered study (https://aspredicted.org/4u7y3.pdf) that will be published in due course. That study investigated whether a purely random competitor would produce the kinds of progressive behavioural changes found here, and could thereby estimate simple time-dependent or exposure-dependent factors.

### Empiricist view versus nativist views of human autonomy

Our work is broadly compatible with an empiricist view of human autonomy as opposed to a nativist view. In our view, some of the key attributes historically associated with “free will”, such as the ability to act endogenously and purposefully, can be acquired, or at least functionally adapted, through experience. Such adaptation requires agents to make novel, non-habitual, ‘smart’ actions appropriate to their current situations. We found that people were more or less successful in adapting their decision biases to boost their performance as the competitive environment changed. An individual’s degree of autonomy is unique and contingent on environmental constraints. A strong nativist view would suggest that autonomy is a state or trait that occurs within the mind of an individual and is independent of the external environment. However, our results imply that autonomous agents are characterised by their capacity to react appropriately to the restrictions that the environment places on their own actions. In this sense, autonomy can be seen as a reasoned, goal-oriented response that occurs through interaction with an environmental context.

## Conclusions

To conclude, we have developed a new experimental paradigm and analysis pipeline to study when and how human actions can become autonomous. We propose a new theoretical construct of *adaptive autonomy*, meaning the capacity to free one’s behavioural choices from constraints of habitual responding, when a particular choice pattern or stereotypical behaviour becomes dysfunctional, for example, due to environmental changes such as the competitive pressure in our game scenarios. We have shown that people can indeed express adaptive autonomy, and that they do so by reducing the elements of their biased action selection patterns, including repetition of choice, rule-based sequential action and dependency on reinforcement. These appear to reflect three distinct forms of adaptive autonomy, rather than a single common cognitive mechanism for avoiding all decision biases, for example by simply strategically boosting the randomness of action choices. We show that becoming free from the effects of reinforcement is particularly difficult. By demonstrating that belief learning plays an important part in adaptive autonomy, we argue for a strong connection between volition, adaptive autonomy and social cognition. Finally, we have looked at behavioural adaptation and cognitive flexibility of action choices through the lens of voluntary action. In *Beyond Freedom and Dignity* (Skinner, 1971/2002) argued that reinforcement-based guidance of conditioned responses obviated the need for any cognitive construct of volition. We have argued a different position. At least in the context of competitive games, the key cognitive features of volition, namely stimulus-independence, habit-independence, and goal-dependence (Haggard, 2019) are precisely what allow flexible, adaptive and successful performance.

## Author’s contributions

Conceptualization, KO and PH; data collection, KO; investigation, KO; formal analysis, KO, LC and PH; Methodology, KO, LC and PH; Software, KO; Project administration, KO and PH; supervision, LC and PH; visualization, KO, LC and PH; writing – original draft, KO; writing – review & editing, KO, LC and PH; funding acquisition, PH.

## Acknowledgements

This publication was made possible through the support of a joint grant from the John Templeton Foundation and the Fetzer Institute. The opinions expressed in this publication are those of the authors and do not necessarily reflect the views of the John Templeton Foundation or the Fetzer Institute. We are grateful to Mr Alexander Platt, Dr Annika Boldt, Dr Stefano Palminteri for their helpful review on the manuscript. We also thank the editor and reviewers for improving our manuscript. We particularly thank Dr Jean Daunizeau for proposing the analyses and models of social cognition.

## Data and code availability

The data frame for a representative participant is available in the supplementary data. The data frames for all the participants and the codes used to generate the figures will be posted when the manuscript is ready for publication after the review process.

