## Supplemental information for "Autonomous behaviour and the limits of human volition"

### Supplementary Methods

#### Exclusion criterion

We encouraged participants to sustain timeout trials under 5%. We checked the histogram of the timeout rates. Seven participants displayed a timeout above 13 %. This was considerably high compared with the other participants (0–5%: 119 participants; 5–8%: 27 participants; 8–11%: 6 participants; 13–20%: 5 participants; >20%: 2 participants). These participants might not be able to follow the instructions or might not be able to keep their attention on the task, thereby being removed from the analysis.

#### Competitor design

The virtual competitor (a flock of birds) was programmed to predict which response interval a participant would select on the current trial. The prediction was made by using the history of the participant's performance up to that trial. Given past behaviour, the competitor predicted which response interval a participant was likely to select. Accordingly, the two other intervals were primed for winning. We adjusted the birds' departure times by taking their travel time (0.25 sec) and the food delivery/travel time (1.5 sec) into account: if a prediction was made on the late interval, the birds departed from the tree at the period of 4.25–5.75 sec, and they reached the delivery point during 4.5–6.0 sec to catch the food when it was delivered.

For programming Competitor 2 and Competitor 3, we looked for the conditional choice probabilities given the past reaction time. In block 2, Competitor 2 sought out sequential patterns – which interval the participant is going to select after the participant made a particular response –. A history of the past 60 trials was used to estimate the conditional probabilities of selecting three intervals given the previous reaction time. The estimated probabilities were conditioned on the last reaction time ± 0.5 sec. Suppose a participant took 2.5 sec to act in the previous trial. Competitor 2 might discover that, in the past, the participant chose the early interval twice, the middle interval twice, and the late interval six times after the participant had acted in 2.0-3.0 sec. In this case, Competitor 2 penalised the late interval 60% of the time. We assumed that using the previous response time (i.e., continuous variable) is more powerful in predicting the next response than using the previous response interval (i.e., categorical variable).

In block 3, Competitor 3 punished outcome dependence – which interval the participant is going to select after the participant made a particular response and won a trial or lost a trial –. A history of the past 60 trials was used to estimate the conditional probabilities of selecting three intervals given the previous reaction time and the previous outcome. Similar to Competitor 2, the estimated probabilities were conditioned on the last reaction time ± 0.5 sec. Given the last reaction time and the last outcome, Competitor 3 looked for the conditional probabilities of three intervals on the upcoming choice.

#### Data analysis

*Quantifying decision bias scores.* Statistical distance is a standardised way of measuring the extent to which the observed probability distribution is different from the target probability distribution. We calculated the Kullback-Leibler divergence to quantify the extent to which the participant's choice probability distribution is different from the choice probability distribution that a bias-free agent would exhibit. See Figure 3.

1) Choice bias. Competitor 1 punished choice preference in selecting one interval more often than the other two. The probabilities of choosing the early, middle and late interval for a bias-free agent would be 0.33, respectively. We computed the choice probabilities $P(c)$ given a history of intervals each participant chose in each block. The K-L divergence is then

$$D_{KL\,choice\,bias} = \sum_{c\,\in\,E,M,L} P(c)\log_2\left(\frac{P(c)}{0.33}\right) \quad (1)$$

2) Transition bias. Competitor 2 punished sequential patterns on the top of choice preferences. Similar to computing the choice probabilities, we computed the conditional probabilities of choosing the early, middle and late interval given the interval chosen on the previous trial $P(c|c_{-1})$ . We measured the K-L divergence of these participant's conditional probabilities from the participant's choice probabilities. The K-L divergence for each previous interval $c_{-1}$ is computed as

$$D_{KL\ c_{-1}} = \sum_{c\,\in\,E,M,L} P(c|c_{-1})\log_2\left(\frac{P(c|c_{-1})}{P(c)}\right) \quad (2)$$

The total K-L divergence as a weighted sum is then

$$D_{KL\,sequential\,bias} = \sum_{c_{-1}\,\in\,E,M,L} P(c_{-1})\cdot D_{KL\ c_{-1}} \quad (3)$$

Since we conditioned the K-L divergence on the previous interval chosen, we took the proportion of observing that situation into account, and we weighed each divergence by this prior probability. The target probabilities (i.e., bias-free agent) were set to be the participant's own choice probabilities, rather than purely random choices 0.33. Therefore, $P(c|c_{-1})$ becomes equivalent to $P(c)$ and the K-L divergence becomes zero, as long as the participant selects three intervals independently from the previous choice (even if the participant favours one interval). This way, we quantified the deviation of patterns associated with the previous choice from sequential patterns logically expected from the participant's own choice probabilities. Competitor 2 specifically detected and punished this conditional dependence.

3) Reinforcement bias. Competitor 3 punished outcome dependence on the top of sequential patterns and choice preferences. Similar to computing the choice probabilities, we computed the conditional probabilities of choosing the early, middle and late interval given the interval chosen and the outcome obtained on the previous trial $P(c|c_{-1}, o_{-1})$. We measured the K-L divergence of these participant's conditional probabilities from the participant's conditional probabilities given the previous interval solely. The K-L divergence for each previous interval $c_{-1}$ and each previous outcome $o_{-1}$ is computed as

$$D_{KL\ c_{-1},o_{-1}} = \sum_{c\,\in\,E,M,L} P(c|c_{-1}, o_{-1}) \log_2\left(\frac{P(c|c_{-1}, o_{-1})}{P(c|c_{-1})}\right) \tag{4}$$

The total K-L divergence as a weighted sum is then

$$D_{KL\,reinforcement} = \sum_{c_{-1}\,\in\,E,M,L} \sum_{o_{-1}\,\in\,\substack{success\\fail}} P(c_{-1}, o_{-1}) \cdot D_{KL\ c_{-1},o_{-1}} \tag{5}$$

Since we conditioned the K-L divergence on the previous interval chosen and the previous outcome obtained, we took the proportion of observing that situation into account, and we weighed each divergence by this joint prior probability. The target probabilities (i.e., bias-free agent) were set to be the participant's own conditional probabilities given the previous interval solely. Therefore, $P(c|c_{-1}, o_{-1})$ becomes equivalent to $P(c|c_{-1})$ and the K-L divergence becomes zero, as long as the participant selects three intervals independently from the previous outcome (even if the participant's choice depends on the previous interval). This way, we quantified the deviation of patterns associated with both the previous choice and the previous outcome from patterns logically expected from the conditional dependence on the previous choice solely. Competitor 3 specifically detected and punished this outcome dependence.

We also quantified the positive reinforcement bias and the negative reinforcement bias, separately (SFig. 1). We computed the K-L divergence of the conditional probabilities given the previous interval and the previous win only or the previous loss only from purely random choices 0.33. Here the statistical distance can be argued as the distance between the participant's post-win behaviour or post-loss behaviour and the bias-free agent who is purely random.

#### Computational models

*Reinforcement learning.* We tested a reinforcement learning (RL) model in which an action value is updated via a Rescorla-Wagner rule (Sutton & Barto, 2018). On each trial, an RL agent selects an action from the early, middle or late interval $a \in E, M, L$. For an action $a$ selected on a trial $t$, the value of action $a$ is updated by a prediction error $\delta$:

$$\delta_t = r_t - V_t(a) \tag{6}$$

where $r_t$ is the actual reward received (1 for successfully avoiding birds and 0 for failure) and $V_t(a)$ is the current expected reward for that action. The reward prediction error $\delta_t$ is then used to update the value of the selected action, weighted by the learning rate $\alpha$

$$V_{t+1}(a) = V_t(a) + \alpha\delta_t \tag{7}$$

*1st order belief learning.* In 1st order belief learning (BL) model, an agent simulates the opponent's intention – which option the opponent is going to select – and decides the action that maximises the expected reward (Camerer, 2003; Hampton et al., 2008; Zhu et al., 2012). Actions $a' \in E, M, L$ are available for the competitor to choose. For each action $a'$ on trial $t$, the player's beliefs about the likelihood of the opponent's action $V'_t(a')$ is updated by a prediction error

$$\delta_t = r_t - V'_t(a') \tag{8}$$

where $r_t$ is 1 for the response interval the opponent has just taken while $r_t$ is 0 for the two other intervals the opponent has not taken. These reward prediction errors are then used to update the likelihoods of the opponent's actions, weighted by the learning rate $\alpha$

$$V'_{t+1}(a') = V'_t(a') + \alpha\delta_t \tag{9}$$

Finally, the agent's values of all three intervals were updated by adding a negative sign to the likelihoods of the opponent's actions

$$V_{t+1}(a) = -V'_{t+1}(a') \quad (10)$$

As the opponent takes the same action interval more frequently, the agent becomes less likely to mimic that interval. See a negative lagged correlation between the opponent's action in preceding trials and the agent's current action (blue line in Figure 5A).

Unlike the RL model, all three action values were updated on every trial because, on a particular trial, the 1st order BL agent observes which option the opponent has taken and which options the opponent has not taken. For this reason, 1st order belief learning is mathematically equivalent to a special case of reinforcement learning model in which action value with the actual reward and action values with hypothetical, fictive rewards are both updated (Abe & Lee, 2011; Camerer & Ho, 1999). In any case, both interpretations involve learning the structure of the game and making use of that knowledge to select the action which maximises the reward.

*2nd order belief learning.* An agent who relies on 2nd order BL simulates the opponent's belief of an agent's action – which option the opponent thinks the agent themselves is going to select (Devaine et al., 2014; Hampton et al., 2008)–. The agent with 2nd order belief assumes that their opponent exploits the 1st order belief. For the agent's action $a$ on trial $t$, the agent assumes that the opponent's beliefs about the likelihood of the agent's action $V'_t(a)$ is updated by a prediction error

$$\delta_t = r_t - V'_t(a) \quad (11)$$

where $r_t = 1$ if the agent themselves has taken the interval $a$. This simulates that the opponent estimates that the agent will repeat that action. For the two other intervals the agent has not taken, $r_t = 0$ was given. The agent with 2nd order belief assumes that their opponent updates the likelihoods of the agent's actions, weighted by the learning rate $\alpha$

$$V'_{t+1}(a) = V'_t(a) + \alpha\delta_t \quad (12)$$

Finally, the agent's values of all three intervals were updated by adding a negative sign to the opponent's action values

$$V_{t+1}(a) = -V'_{t+1}(a) \quad (13)$$

As the agent takes the same action interval more frequently, the opponent becomes more likely to intercept that interval. Therefore, the agent themselves becomes less likely to repeat the same interval. See a negative lagged correlation between the agent's preceding actions and their current action (green line in Figure 5A).

*Modelling choice probabilities.* For all models, the agent's action values were converted into the choice probabilities using the soft-max function to simulate action selection,

$$P_t(a) = \frac{e^{\beta\cdot(V_t(a)+b(a))}}{\sum_{a\in E,M,L} e^{\beta\cdot(V_t(a)+b(a))}} \tag{14}$$

where $P_t(a)$ is the probability of choosing the interval $a$. The inverse temperature parameter $\beta$ scales the relative difference between the choice probabilities, which scales decision uncertainty. We added the decision preference term $b$ with an exponential temporal discounting (Story et al., 2014):

$$b(a) = e^{-\rho\cdot T(a)} \tag{15}$$

where $T$ is the time corresponding to the chosen interval ($T = 0$, 1.5 or 3.0 sec for the early, middle or late interval, respectively). The parameter $\rho$ scales the relative preference to earlier intervals, which captures an individual's temporal discounting or an individual's trend to respond immediately.

*Model fitting and evaluation.* For each model, we fitted the model choice probabilities to the participant's choice interval data by minimizing the negative log-likelihood of the observed choices using Bayesian adaptive direct search (BADS) (Acerbi & Ma, 2017). Free parameters were optimised individually for each participant and separately for each block with the following boundaries: $\alpha \in [0, 1]$, $\beta \in [0, 20]$, $\rho \in [0, 0.2]$. For the stochastic action generation model, we fixed the parameter $\alpha$ to 0 so that this model could only capture the participants' decision uncertainty and their choice preference. Across all trials, this model produced constant model choice probabilities with preference for earlier intervals. In this nested structure, a better model index relative to the stochastic action generation model would indicate the presence of an update process. To verify that we had found the global minimum, we repeated the search process with different starting points. For model comparison, we applied AICc—Akaike information criterion with a correction for finite sample size—to each participant and model as the information criterion for goodness-of-fit (Burnham & Anderson, 1998; Hurvich & Tsai, 1989). The formula for AICc adds an extra penalty term $\frac{2K^2+2K}{N-K-1}$ to the formula for AIC by taking into account the number of model parameters $K$ and the number of data points $N$. This penalty avoids potential overfitting and helps select the models that have fewer parameters as Bayesian information criterion (BIC) does. Based on AICc values across participants and across models, we computed the protected exceedance probabilities (Rigoux et al., 2014;

Stephan et al., 2009). See Figure 5D. The parameter estimates best fitted by each class of the model are illustrated in SFigure 2.

*Simulating model agents.* To compute lagged correlations in model agents (Figure 5A), we simulated that an agent exploits either a stochastic action generation, reinforcement learning, 1st order belief learning or 2nd order belief learning and plays against the prediction algorithm used in the actual experiment. The competitor's prediction was made using a history of choices made by each class of model agent. For each iteration of 60 trials (which is approximately equal to the number of trials in the actual experiment), we generated a sequence of agent's action choices and a sequence of opponent's action choices. We iterated this step 10,000 times and computed lagged correlations with the current choice. We verified that changing model parameters affects the coefficients of lagged correlations but the pattern of correlation (e.g., a negative lagged correlation with own preceding choices in 2nd order belief learning) remains the same. We also verified that the pattern of lagged correlation remains the same regardless of different competitor algorithms. In Figure 5A, we illustrate the lagged correlations in the following set of parameters (for the stochastic model, $\alpha = 0$, $\beta = 0$, $\rho = 0$; for the other three models, $\alpha = 0.3$, $\beta = 5$, $\rho = 0$) and when model agents played against Competitor 1 who predicts the choice bias.

### Supplementary Results

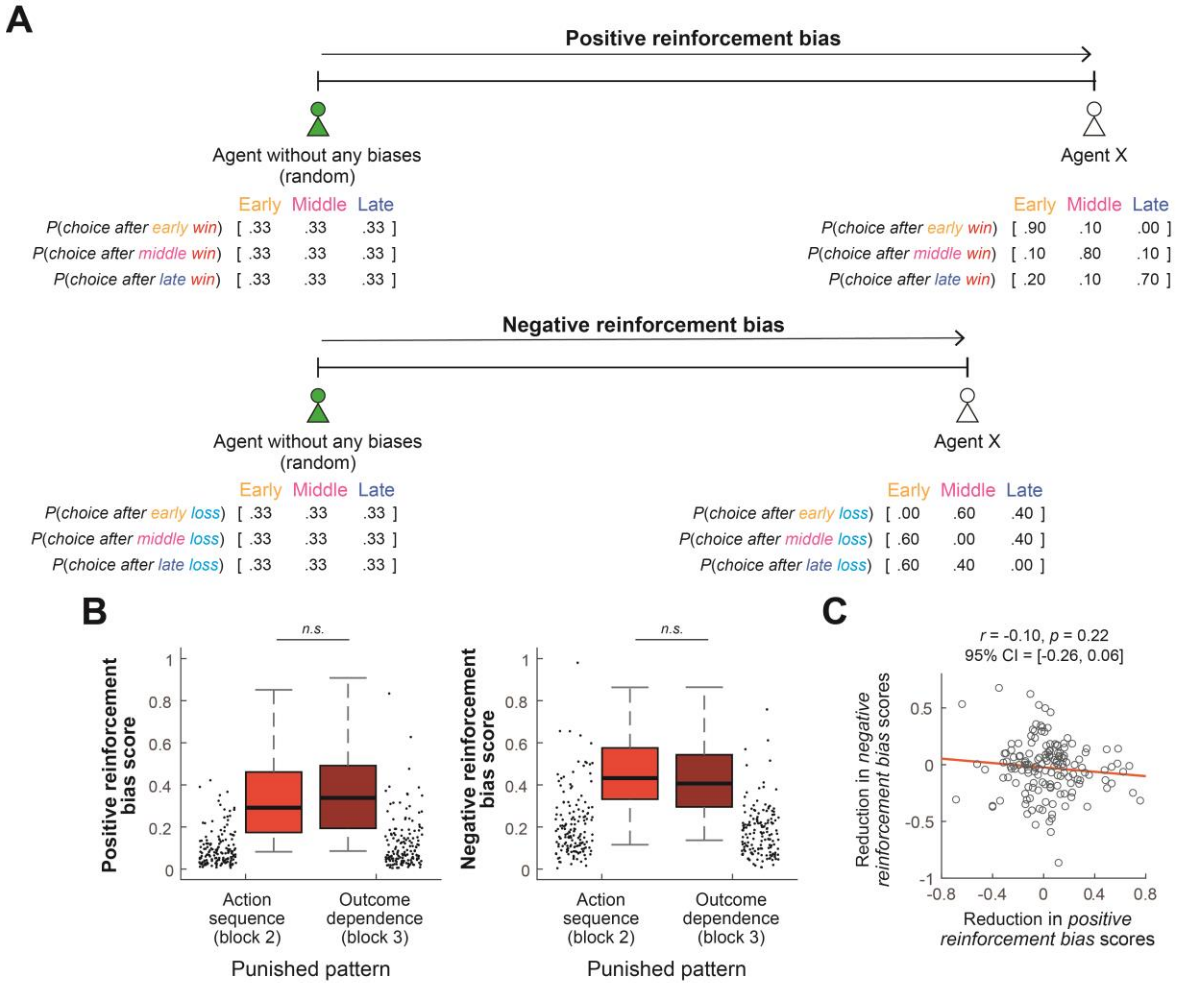

**SFigure 1. Positive and negative reinforcement bias**

**A.** The reinforcement bias vector in Figure 3C is broken down into positive reinforcement bias and negative reinforcement bias. The statistical distance between an agent without any biases (purely random agent) and participants is computed. **B.** Planned comparisons show the degree of adaptive autonomy (bias reduction) when outcome dependence is punished. **C.** There is no correlation between the bias reduction in the positive reinforcement bias and the bias reduction in the negative reinforcement bias. The ability to adapt away from a win-stay type behaviour is not associated with the ability to adapt away from a lose-shift type behaviour across participants.

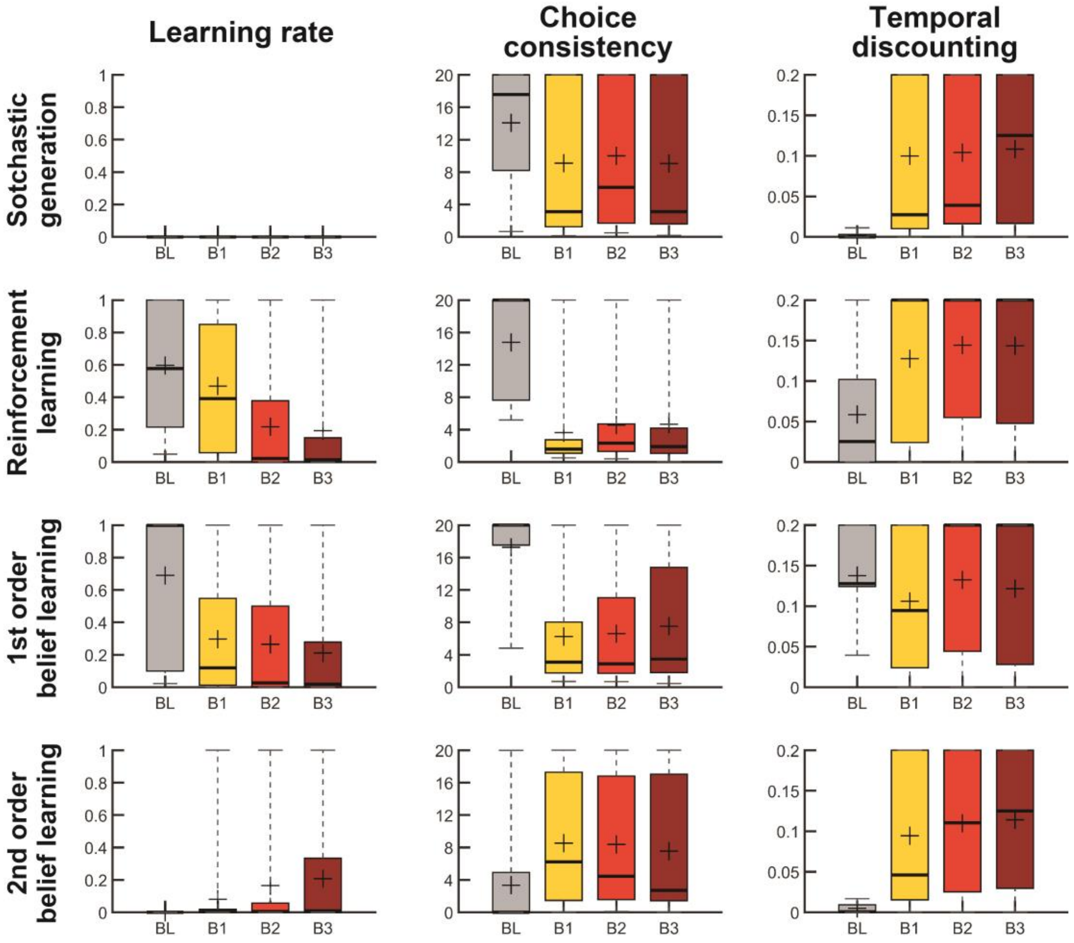

**SFigure 2. Parameters of the model fit**

We show the parameter estimates best fitted by each class of the model across blocks. Learning rate, $\alpha$: The larger the value, the quicker the update. Choice consistency, $\beta$: The larger the value, the more consistent in choosing a higher value option. Temporal discounting, $\rho$: The larger the value, the more likely it is to choose earlier intervals. On each box, the central horizontal mark represents the median, the cross mark represents the mean, the edges of the box are the 25th and 75th percentiles and the whiskers are the 2.5th and 97.5th percentiles.
